# Characterization of Quantum Dots with Hyperspectral Fluorescence Microscopy for Multiplexed Optical Imaging of Biomolecules

**DOI:** 10.1101/2022.03.17.484752

**Authors:** Shuyan Zhang, Joseph Yong Xin Cheng, Jian Jun Chua, Malini Olivo

## Abstract

The optical properties of quantum dots were extensively characterized using a hyperspectral fluorescence microscopy system. The system provides a single excitation wavelength in the ultraviolet and 311 emission wavelength channels in the visible. This allows detection of multiple fluorophores (e.g. different quantum dots) with a high spectral resolution in one go which is not achievable with a conventional fluorescence microscope where different filter sets have to be used. A spectral library was established based on the spectral profiles of six types of quantum dots. Notably, a slight spectral shift was observed for all samples while the sample was drying. Subsequently, two quantum dot mixture samples were studied. Using the spectral unmixing approach, the relative proportions of each quantum dot within a homogeneous mixture and the spatial distribution of each quantum dot within a heterogeneous mixture were calculated. The calculated values match well with the theoretical predictions. Hence, the analysis method presented here can be used for simultaneous imaging of multiple fluorophores using hyperspectral imaging technology. The results provide valuable information for the realization of real-time multi-channel in vivo fluorescent imaging of biomolecules.

## INTRODUCTION

Quantum dots (QDs) are nanometer-sized crystals, typically of semiconductor material and ranging from 2 - 10 nanometers in diameter, which exhibit fluorescence: a physical phenomenon in which a substance is excited by photons of a shorter wavelength and subsequently releases photons of a longer wavelength as it relaxes back into its ground state. ^1^ The small size of quantum dots leads to a discretization of the energy levels within the semiconductor material and gives rise to the size-dependent fluorescence spectra of quantum dots: a decrease in the crystal size results in a larger energy bandgap and thus blueshifts the excitation and emission spectra of the quantum dot. ^2^ Apart from the size tunability of their fluorescence spectra, other key features of quantum dots that make them highly attractive fluorophores include high photoluminescence quantum yield, ^3^ broad excitation coupled with narrow and symmetrical emission spectra, ^4^ high resistance to photobleaching, ^5^ and ease of surface modification. ^6^

Since their discovery, quantum dots have been utilized in various bioimaging applications; ^7^ such as cellular labeling, ^8–10^ invivo tissue imaging, ^11–13^ bioassays, ^14–17^ and as Förster Resonance Energy Transfer (FRET) donors; ^18–20^ where their high quantum efficiency and high photostability afford good detection sensitivity and allow for imaging to be performed over extended periods of time. Owing to their well-studied surface chemistries, ^21^ quantum dots have been conjugated to a variety of biomolecules, including antibodies, ^15,22^ proteins, ^23–25^ and oligonucleotides; ^16,17,26^ to achieve specific targeting of various biomarkers of interest. Additionally, given the narrow and symmetrical emission spectra of quantum dots which minimize their spectral overlap with other fluorophores, as well as their broad excitation spectra which allows for multiple quantum dots to be excited from a single UV wavelength source; quantum dots possess great potential towards multiplexed imaging and have seen repeated use in such applications. ^14,15,17,27,28^ Xu et al. utilized multiple quantum dot-antibody conjugates to perform simultaneous immuno-histo-fluorescence (IHF) staining of different targets in formalin-fixed, paraffin-embedded (FFPE) tissue slides of human head and neck cancer. ^14^ Peng et al. studied the patterns of tumor invasion in human gastric and breast cancer FFPE tissue slides via multiplexed IHF. ^28^ Chan et al. utilized multiple quantum dot-labeled oligonucleotide probes for fluorescence in situ hybridization (FISH) imaging of mRNA biomarkers in mouse brain sections. ^17^

The technique of utilizing fluorophores such as quantum dots to label and image particular features in a sample is known as fluorescence imaging, ^29^ on the basis that fluorophores can provide better visual contrast and localization information for the features of interest as compared to an unlabeled sample. Currently, there are two broad types of microscopes used in fluorescence imaging. A widefield fluorescence microscope typically uses a broadband incoherent light source such as a light emitting diode (LED), and a pair of excitation and emission filters which serve the purpose of selecting the desired excitation wavelength and filtering out wavelengths other than the fluorescence emission, respectively. ^30^ On the other hand, a confocal laser-scanning fluorescence microscope utilizes a high-intensity monochromatic light source, such as a laser, which determines the excitation wavelength, while the emission wavelengths are selected by emission filters. ^31^ The high spatial resolution and contrast in the acquired images, due to the optical sectioning capability which confocal microscopes are well known for, is achieved by the use of a spatial pinhole to block out-of-focus light from other focal planes in the sample.

While fluorescence imaging can acquire spatial information regarding the localization of fluorophores across a sample, it cannot identify each fluorophore according to its unique spectral fingerprint; for this to be accomplished, fluorescence spectroscopy has to be performed instead. Conversely, while it is able to acquire spectral information to deduce the type of a particular fluorophore and its intensity, the technique of fluorescence spectroscopy is only able to do so for a single point within a sample and lacks the ability to visualize the localization of fluorophores across the sample. Currently, emission filters are used to isolate the spectral band corresponding to the fluorescence emission of each fluorophore, thus allowing for one fluorophore to be imaged at a time; as such it is not strictly necessary for fluorophores to be identified by their spectral fingerprint. However, imaging multiple fluorophores with different spectral characteristics remains a tedious process, requiring the use of separate excitation and emission filters that are specifically optimized for each fluorophore, which has to be swapped out before imaging each fluorophore; as such the sample has to be imaged sequentially for each fluorophore used. Unconventional fluorophores thus pose an added challenge when they do not match well with the spectral bandwidth of commonly used emission filters. Furthermore, it is especially challenging to identify fluorophores with highly similar emission spectra and large spectral overlap. This may impose limitations on the experimental design when multiple fluorophores are intended to be used simultaneously or when imaging targets in the sample are located close to each other spatially.

Hyperspectral imaging (HSI) is an imaging technique that combines the principles of optical imaging with spectroscopy, capturing three-dimensional images comprised of information from the spatial dimensions (x, y) and the spectral dimension (λ). ^32^ As such, each pixel in a hyperspectral image contains the complete spectrum of the sample imaged by that pixel, and conversely, each wavelength on the spectrum contains its corresponding single-channel image. This technique is useful in multiplexed fluorescence imaging where each fluorophore within the sample can be identified by its spectral characteristics and at the same time, the spatial localization of each type of fluorophore can be elucidated. Two recent review articles from Fakhrullin’s team and Rivera_Gil’s team discussed the use of hyperspectral imaging combined with darkfield imaging modality for detecting nanoscale particles such as quantum dots for environmental and biological research. ^33,34^

In this paper, we present a customized wide-field hyperspectral fluorescence microscopy system based on a liquid crystal tunable filter that circumvents the challenges faced in fluorescence imaging for multiplexed detection. Compared to other fluorescence microscopy systems, our system does not rely on preset excitation and emission filters to excite and acquire fluorescence emissions from a sample, which limits these other systems from simultaneously acquiring fluorescence signals from multiple different fluorophores and makes multiplexed imaging a tedious process. Instead, our system features a liquid crystal tunable filter paired with a monochromatic complementary metal oxide semiconductor (CMOS) camera to sequentially acquire grayscale images at each spectral channel (up to 1 nm step size) within the visible wavelength scanning range in a spectral scanning format, which are then stacked together to form a three-dimensional hyperspectral image. Compared to angle-tuned filters or filter wheels, the liquid crystal tunable filter contains no moving parts, thus the optical pathway through the system is not affected during operation and the captured images are not subject to any undesirable pixel shift effect. The liquid crystal tunable filter can achieve a wavelength switching speed of less than 200 ms. Here, we demonstrate the capabilities of our microscope for multiplexed fluorescence imaging and present a systematic way of studying the optical properties of fluorophores based on their spectral features. We characterized the fluorescence spectra of a series of six types of quantum dots with different emission peaks and discovered their spectral differences in wet and dry conditions. We also studied a homogenous mixture and a heterogeneous mixture of quantum dots. We showed how to use the spectral unmixing technique to identify each quantum dot type, quantify its relative proportion, and elucidate its spatial localization within the mixture sample. Lastly, we proposed a multiplexed sensing framework for quantum dot-labeled biomolecules using hyperspectral fluorescence microscopy. The results here provide valuable information for realtime simultaneous multi-channel fluorescence imaging in the future.

## METHODOLOGY

### Hyperspectral Fluorescence Microscopy System

The customized hyperspectral fluorescence microscopy system mainly consists of a UV LED light source (M375L4, Thorlabs), a liquid crystal tunable filter (Kurios-VB1, Thorlabs), and a monochromatic CMOS camera (CS126MU, Thorlabs). Figure 1(a) shows a schematic of the optical pathway of the system. The light source outputs at the central wavelength of 375 nm with a typical output power of 1540 mW, which is sufficient to excite all six types of quantum dot samples, given the broad excitation spectra of our samples. The incoming light passes through a collimation lens to produce collimated light to a beam splitter, which is then focused onto the sample by an objective lens. The scattered light from the fluorescence emission of the sample is collected by the same objective lens and passes through the beam splitter before reaching the liquid crystal tunable filter, which isolates the light of a narrow band of wavelengths corresponding to the desired spectral channel. The tunable filter covers a range of 420 - 730 nm with a minimum step size of 1 nm and a bandwidth of 18 nm for each spectral channel, resulting in a maximum possible 311 spectral channels that can be captured; implying that in theory, a total of 311 different types of fluorophores can be potentially identified and their spatial distribution elucidated. Each single-channel image is then captured by the CMOS camera, which captures 12-bit grayscale images of a high resolution of up to 4096 × 3000 pixels at a maximum frame rate of 21.7 fps, with a pixel size of 3.45 μm × 3.45 μm. Finally, all single-channel images from each individual scan wavelength are stacked together to form a hyperspectral image.

**Figure 1.**
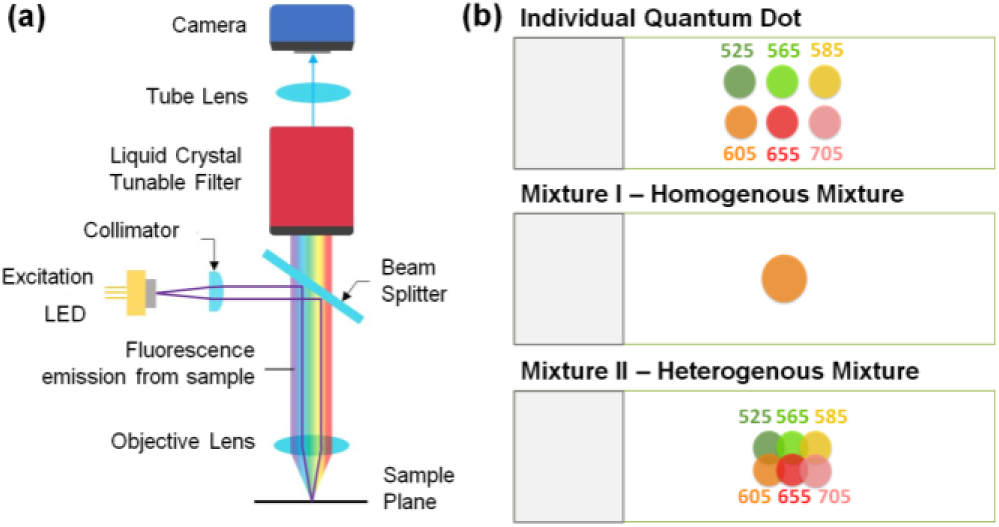
(a) Optical pathway for fluorescence imaging. (b) An example of the sample, i.e. layout of mixture (II) of all six quantum dot samples.

### Sample Preparation

The quantum dot samples used in this paper were CdSe core, ZnS shell structure (CdSe/ZnS) quantum dots coated with an additional polymer shell and conjugated to streptavidin protein (Q10151MP, Invitrogen). A total of six quantum dot conjugates with different emission spectra were used, with emission peaks at 525 nm (Qdot 525), 565 nm (Qdot 565), 585 nm (Qdot 585), 605 nm (Qdot 605), 655 nm (Qdot 655), and 705 nm (Qdot 705) respectively. All samples were obtained as 1 μM solutions in 1 M betaine and 50 mM borate solvent with 0.05% sodium azide at pH 8.3. For imaging individual quantum dots, 2 μL of each sample was loaded onto a clean glass slide, while for imaging quantum dot mixtures, two types of mixture samples were prepared.

Mixture (I) was a homogenous mixture of all six quantum dot samples. 0.5 μL of each quantum dot conjugate was added and mixed using a vortex mixer, yielding a total volume of 3 μL of the mixture; out of which, 2 μL of the mixture was used for imaging.

Mixture (II) was a regionally separated mixture of all six quantum dot samples, prepared by loading 1 μL of Qdot 525 and Qdot 565 each, and 0.5 μL of the remaining samples onto a clean glass slide without prior mixing. As shown in Figure 1(b), samples were loaded into a 3 by 2 matrix format on the glass slide. As samples were loaded close to each other, there were overlapping regions containing heterogeneous mixtures of two different samples; for instance between Qdots 525 and 605, as well as between Qdots 585 and 705.

All individual samples and mixture (I) were imaged while still wet as well as after drying, while mixture (II) was imaged only after drying.

### Image Acquisition Process

Hyperspectral images for all samples were captured with the centermost 1000 × 1000 pixels of the sensor array, except for mixture (II) which was captured with the centermost 1500 × 1500 pixels; using a scan range of 420 nm to 730 nm with a spectral step size of 1 nm, for a total of 311 spectral channels acquired in each image.

For imaging of individual quantum dot samples and mixture (I), hyperspectral images were acquired under 10x magnification. The exposure duration of 170 ms for all individual samples and mixture (I) was selected based on the emission of the brightest sample, Qdot 655, such that the hyperspectral image of Qdot 655 would be as bright as possible without any overexposure. The identical exposure settings used across all aforementioned samples allow for a fair comparison between the peak intensities of the various quantum dots. Mixture (II) was imaged under 4x magnification to preserve a sufficiently wide field of view during image capture. An exposure duration of 900 ms was selected for optimal exposure of the different regions of interest across the sample, which were the overlapping regions between different quantum dots.

After the acquisition of the hyperspectral images, full-color reconstruction was performed by extracting and merging three single-channel images corresponding to the Red (633 nm), Green (545 nm), and Blue (455 nm) channels; followed by contrast enhancement via linear stretching on the RGB image. This allows the hyperspectral image to be quickly visualized and gives an overview of the spatial information captured within the hyperspectral image.

### Spectral Unmixing

Spectral unmixing is an analysis tool that studies the spectrum of a sample containing multiple substances by decomposing the overall spectrum into the set of its component spectra, or endmembers; as well as their respective proportions, or abundances. ^35^ For hyperspectral imaging, spectral unmixing can be used in situations where each pixel in a hyperspectral image contains information from a mixture of multiple substances at the sub-pixel scale, or when imaged substances may possess similar spectra and result in high degrees of spectral overlap with each other; in both cases manifesting as a superposition of the spectra of multiple components in the observed spectrum at each image pixel.

While there are many mathematical models describing various types of interactions between the different substances at the sub-pixel scale, the linear spectral unmixing model is the simplest and is suitable for describing substances that do not interact with each other. Each photon that reaches the detector is assumed to have only interacted with a single substance with no other optical effects along its pathway to the detector. Mathematically, linear spectral unmixing for *N* different endmembers can be expressed as the following equation:

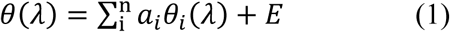

where *θ*(*λ*) is the superimposed spectrum as a function of the wavelength, *a*_*i*_ and *θ*_*i*_(*λ*) are the abundance and endmember spectrum for the *i*-th endmember respectively, and *E* refers to the observation noise term. Typically, endmember extraction is first performed to determine the set of endmember spectra that comprises the superimposed spectrum. Once the set of endmember spectra (i.e. a spectral library) is known, the abundances of each endmember can then be estimated by the leastsquares solution to equation (1), typically with non-negative and sum-to-one constraints set upon the estimated abundance values if all endmembers are already known beforehand.

In this paper, linear spectral unmixing was performed on the hyperspectral images of mixtures (I) and (II). The set of endmember spectra for these images was the spectra of individual quantum dots obtained from the hyperspectral images of the individual samples; as such, endmember extraction does not have to be conducted. The unmixing process was performed on each pixel in the hyperspectral image, taking the superimposed spectrum as well as the known endmember spectra as inputs, and returning the series of calculated abundance values of each endmember as the output. Thus, spectral unmixing on hyperspectral images yields a new data cube of identical spatial dimensions and a reduced number of spectral channels corresponding to the number of endmembers, with each pixel containing the series of unmixed abundance values from its original spectrum.

## RESULTS AND DISCUSSION

Unlike conventional fluorescence imaging, hyperspectral fluorescence imaging allows for the quantification of the relative emission intensities of each quantum dot sample present within each pixel in the case of mixture (I), and the visualization of the spatial distribution of each quantum dot sample across each image in the case of mixture (II), by performing spectral unmixing on the spectral information obtained from the hyperspectral images.

### Individual Quantum Dot Samples

For each sample, we took three hyperspectral images when the sample was wet and one hyperspectral image after the sample had dried. Figure 2(a) shows the full-color images reconstructed from the hyperspectral images of the edges of the six quantum dot samples when the samples were dry, with the “coffee-ring” effect visible in these images. Figure 2(b) shows the emission spectrum of each sample, which was generated from the hyperspectral images of the wet samples (see Supporting Information, Figure S1) since the samples were more uniform in the wet condition. The average emission spectrum was calculated across the entire field of view for each hyperspectral image. The obtained spectra closely matched with manufacturer-provided data in terms of the shape, position, and width of the emission peaks; thus demonstrating that our hyperspectral fluorescence microscope system is able to capture the spectral information of imaged samples with a high degree of accuracy. The obtained spectra were subsequently used to build the spectral library for the spectral unmixing analysis of mixture (I) and mixture (II). It should be noted that the average spectra of the individual quantum dot samples had different peak intensities, due to differences in the extinction coefficients of each quantum dot sample for the same excitation wavelength and intensity.

**Figure 2.**
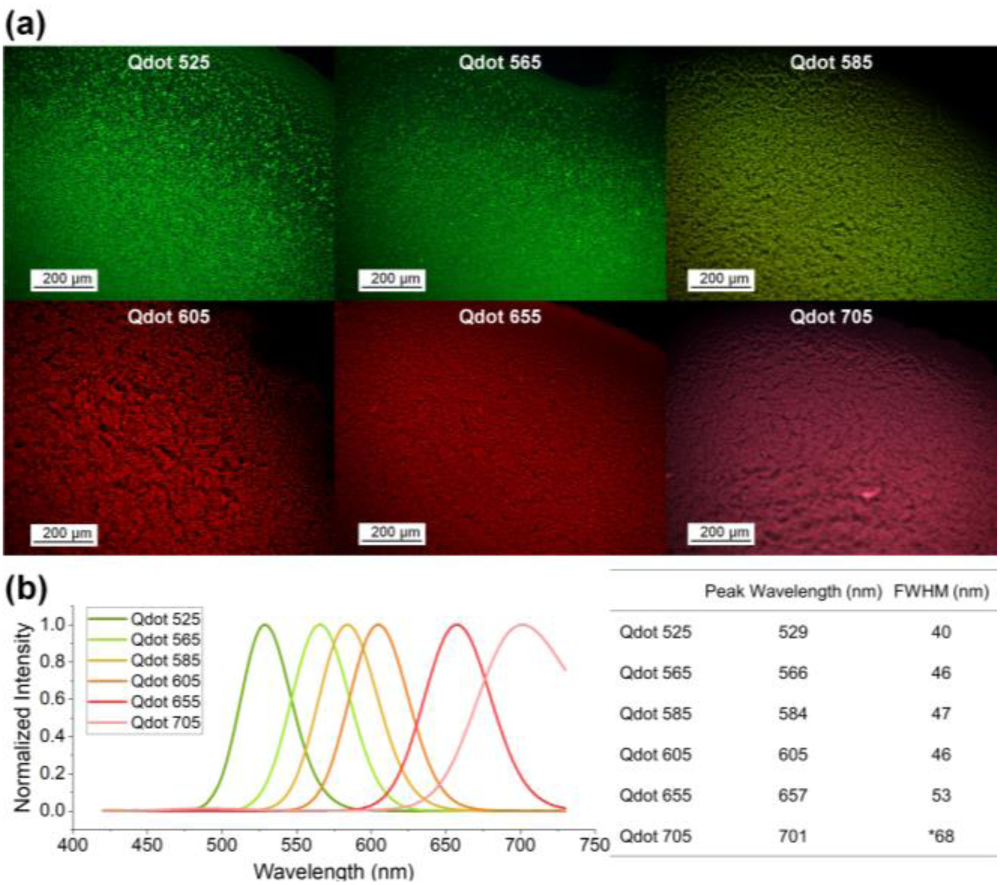
(a) Reconstructed full-color images of individual quantum dot samples (edge of sample, 10x magnification) after samples had dried. Images on display were captured using a larger area of the sensor array (3200 × 2500 pixels) and with a larger spectral step size of 5 nm. Note that these were not used for building the spectral library. (b) Normalized average emission spectra of individual quantum dot samples with peak wavelengths and FWHM values shown. The peak intensities of all samples were scaled to the same value for clearer representation. *The exact FWHM value of the Qdot 705 sample could not be determined as the full emission profile was out of the spectral scanning range of the tunable filter; hence an estimated value is shown assuming that the spectral profile is symmetrical.

Comparing the hyperspectral images of the same region of the same sample in the wet and dry conditions, as shown in Figures 3(a) and 3(b), respectively. Interestingly, we observed that there was a spectral redshift of around 5 nm, as shown in Figure 3(c). This spectral shift was observed in all of the samples (see Supporting Information, Figure S2), which is likely due to the aggregation of the quantum dot particles during the drying process over a period of time (approximately 15 minutes in our case). ^36^ We note that this has not been discussed before in the literature and it may have a wide implication for nanoparticles-based fluorescent imaging.

**Figure 3.**
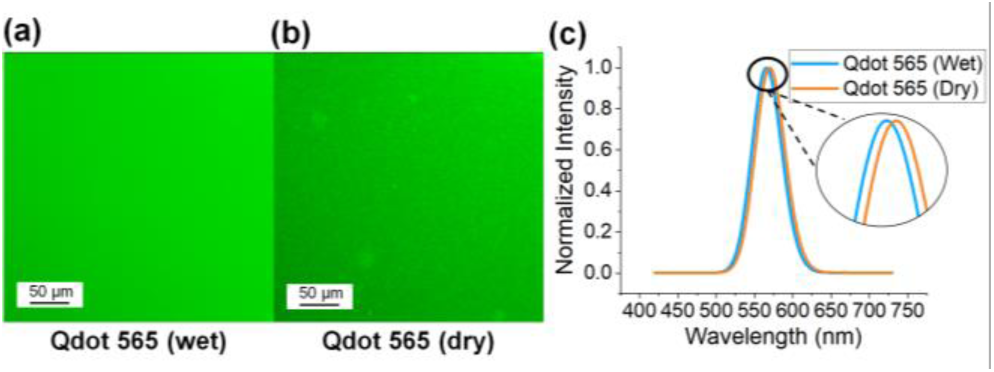
(a) Reconstructed full-color image of Qdot 565 sample when wet and (b) dry. (c) Normalized average emission spectra from the hyperspectral images of (a) and (b), with a peak intensity shift of around 5 nm.

### Mixture (I) - Homogeneous Mixture of Quantum Dot Samples

To evaluate the multiplexed detection capabilities of the system, mixture (I) containing a homogeneous mixture of equal proportions of all six quantum dot samples was imaged. Linear spectral unmixing was performed on the emission spectra of mixture (I) with six endmembers; each endmember corresponding to the emission spectrum of each quantum dot sample as obtained in the previous section. This allows us to calculate their relative proportions in the mixture solution.

Figure 4 shows the reconstructed full-color images of the same region of mixture (I) while both wet and dry, as well as the average spectra and spectrally unmixed proportions of each type of quantum dot sample within the hyperspectral images. Figure 4(a) displays a uniform orange color across the field of view, while Figure 4(d) displays an overall red color with patches of uneven quantum dot aggregates.

**Figure 4.**
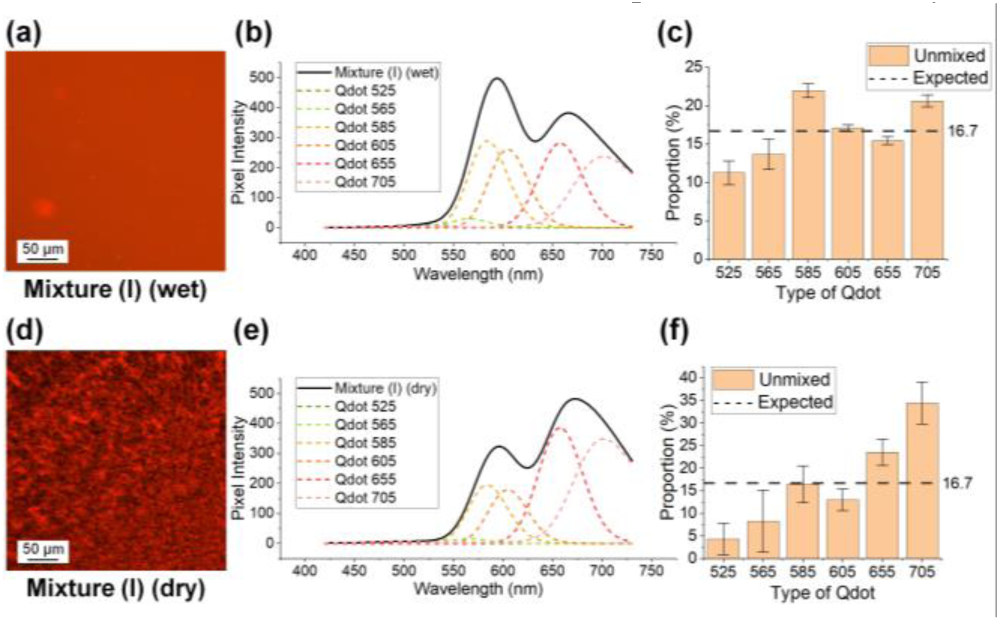
(a) Reconstructed full-color image of the mixture (I) of all six quantum dot samples (center of the sample, 10x magnification) in the wet condition. (b) Measured emission spectrum (solid line) and decomposed spectra (dotted lines) of (a). (c) Spectrally unmixed proportions of each type of quantum dot sample of (a). (d) A reconstructed full-color image of the mixture (I) in the dry condition. (e) Measured emission spectrum and decomposed spectra of (d). (f) Spectrally unmixed proportions of each type of quantum dot sample of (d).

Figure 4(b) and (e) show the emission spectra of mixture (I) in the wet and dry conditions, respectively. The solid lines are the observed superimposed spectrum of the mixture and the dotted lines are the decomposed spectra of the corresponding endmembers. Both the wet and dry samples appeared to have two prominent peaks at approximately 595 nm and 670 nm, with the fluorescence emissions of Qdot 585 and Qdot 605 contributing to the peak at 595 nm and the fluorescence emissions of Qdot 655 and Qdot 705 contributing to the peak at 670 nm.

As the fluorescence emissions of Qdot 525 and Qdot 565 were much weaker compared to the rest of the quantum dot samples for the same excitation condition, their contributions were much less significant. Unlike the individual quantum dot samples, there was a noticeable change in the shape of the spectra between the wet and dry samples of mixture (I) which explains the differences in color of their respective full-color images; with the 595 nm peak having a higher intensity than the 670 nm peak in the wet sample, while the 670 nm peak having a higher intensity than the 595 nm peak in the dry sample. Two possible factors that could contribute to the spectral shift between the wet and dry samples are the aggregation of quantum dot particles as mentioned previously, as well as the inter-quantum dot energy transfer from the quantum dots with lower peak wavelength to those with higher peak wavelength, thus quenching the emission intensity of the quantum dots with a lower peak wavelength while strengthening the emission intensity of the quantum dots with a higher peak wavelength. ^37^ This inter-quantum dot energy transfer is reflected in the differences between the unmixed proportions of the wet and dry sample, with the dry sample displaying a general linearly increasing trend with the peak wavelength of the quantum dot samples.

Figure 4(c) and (f) are the expected and calculated (spectrally unmixed) relative proportions of each quantum dot sample in the mixture in the wet and dry conditions, respectively. The expected proportions are shown as dashed lines and the calculated or unmixed proportions are shown as bar graphs. The relative proportions were calculated based on the decomposed peak intensities of each quantum dot sample since there is a linear relationship between the peak intensity and the concentration of the sample (see Supporting Information, Figure S3). The calculated proportions are relatively close to the expected values with the wet sample matching better than the dry sample (see Supporting Information, Figure S4). The discrepancies between the calculated proportions and the expected values are likely due to the interactions between different quantum dot samples, which suggest that a nonlinear spectral unmixing may be a more accurate representation of the mixture sample. ^38^ Nonetheless, the linear spectral unmixing model adopted here is simple enough to be used for estimated quantitative analysis.

### Mixture (II) - Heterogeneous Mixture of Quantum Dot Samples

In mixture (I), we focused on the spectral features of the hyperspectral images. Hence, we prepared mixture (II) which was a heterogeneous mixture of the quantum dots to study the spatial features. We demonstrate our system’s ability to spatially resolve different fluorophores in a given sample. As shown in Figure 1(b), mixture (II) contains multiple regions where two quantum dot samples overlap spatially to form a heterogeneous mixture.

Figure 5(a) shows the reconstructed full-color image of the mixture sample. Subsequently, spectral unmixing was performed on the hyperspectral images to spatially distinguish the different regions in each image from each other based on the constituent quantum dot sample. Since spectral information is saved at each pixel, an image can be processed to show regions where the spectral features are the same or similar. For each imaged region, only the quantum dot samples present within the region were included in the unmixing process as endmembers. Figure 5(b) and (c) show the spatial distribution of each quantum dot sample based on the spectral unmixing results as a grayscale image with brighter pixels indicating a higher concentration. Note that the spatial distributions obtained using this method are more accurate than those obtained based on the single-channel images at the peak emission wavelength of each quantum dot sample because spectral unmixing takes into account the whole spectral information and the spectral overlap between different quantum dots in each spectral channel.

**Figure 5.**
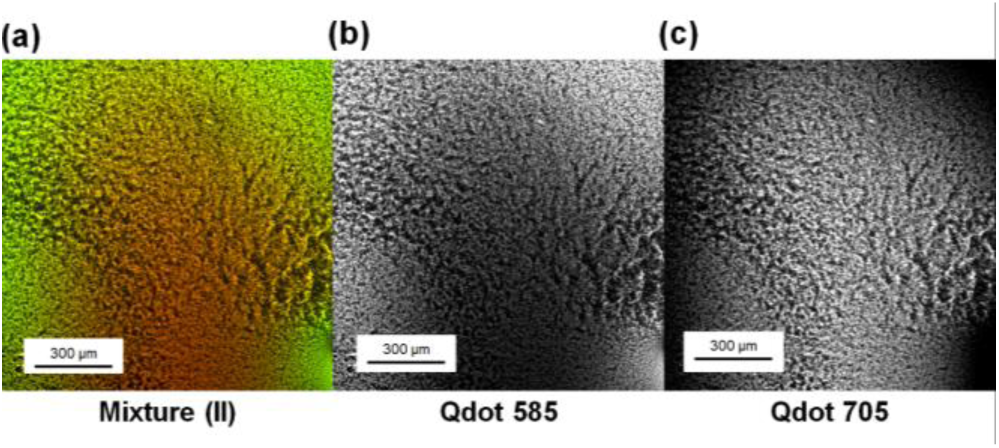
(a) Reconstructed full-color image of the region of overlap between Qdot 585 and Qdot 705 (4x magnification). (b) Spectrally unmixed grayscale images displaying the spatial distributions of Qdot 585 and (c) Qdot 705.

### Framework for Multiplexed Imaging

Given the favorable optical properties of quantum dots as fluorophores and the fact that different quantum dots can be excited at the same wavelength (i.e. same light source), quantum dots are very well suited for multiplexed imaging. Here, we propose a framework for multiplexed imaging where different quantum dots are used as fluorescent labels for different target biomarkers in a sample. We have shown that our fluorescent hyperspectral imaging system can resolve the spectral and spatial information with high resolution at one go without the need to change filters or move stages. Compared to a more conventional approach where the sample is sequentially stained and imaged for each target biomarker, our framework allows for staining to be performed for all target biomarkers and only a single round of imaging is needed for the entire sample.

As shown in Figure 6(a), quantum dots are first conjugated to biomolecules capable of binding to the target biomarker through specific biomolecular interactions (e.g. antibodies binding with antigens or complementary oligonucleotide sequences) to form sets of quantum dot probes, with different sets targeting at different biomarkers. The sample is then sequentially stained with the various sets of quantum dot probes, adding each set of probes without removing the previous set, such that the sample is eventually stained with all sets of probes at the same time. Finally, the stained sample can be imaged in a single step using our system. Spectral unmixing can then be performed to isolate the signals corresponding to each target biomarker, such that (1) the relative proportions of each biomarker at each image pixel can be compared and (2) regions containing each target biomarker can be viewed individually, as demonstrated in the previous sections. When the number of target biomarkers becomes large and the different types of quantum dots are not sufficient for labeling all target biomarkers, one can also use different intensity ratios of the quantum dots as an added dimension for optical coding, as shown in Figure 6(b). Some pioneering work was done by Han et al. and Liu et al. where quantum dot-tagged microbeads were used for targeting specific DNA oligonucleotide sequences. ^27,39^

**Figure 6.**
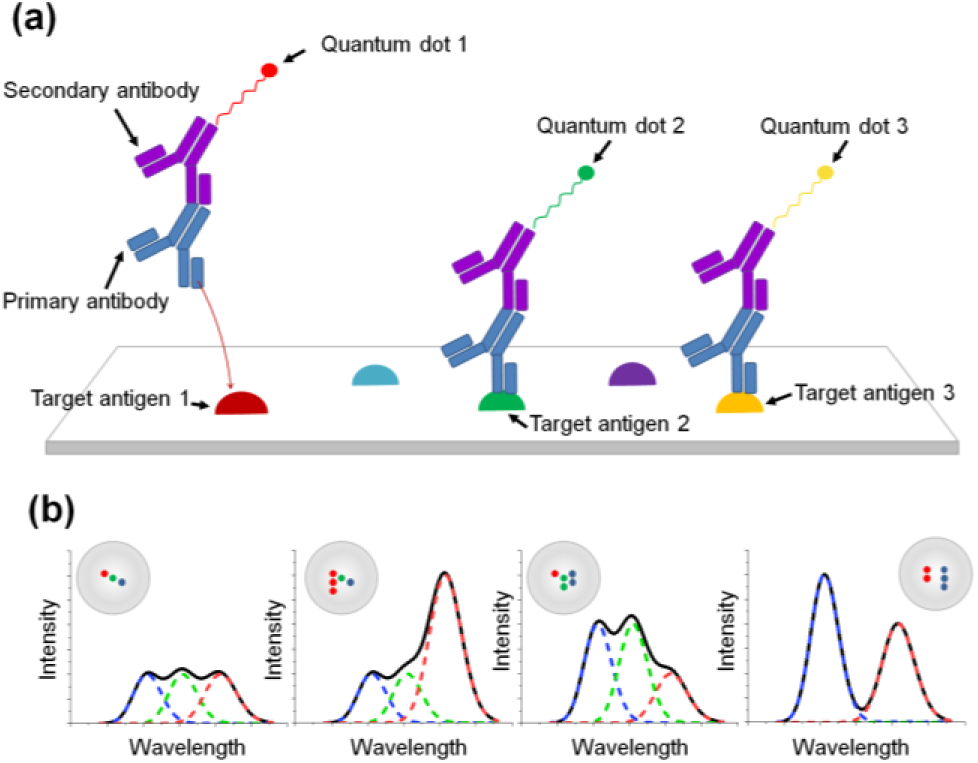
(a) Illustration of multiplexed staining with three different quantum dot-antibody conjugates. (b) Illustration of optical coding using quantum dot-tagged microbeads.

## CONCLUSION

We have presented a widefield hyperspectral fluorescence microscopy system that is able to capture both spatial and spectral information of fluorophores with high resolution and accuracy. We have demonstrated its capabilities by characterizing a series of six different types of quantum dot samples with different emission peaks, as well as two mixtures, one homogeneous and one heterogeneous mixture, of all six types of quantum dots. From the hyperspectral images of the individual quantum dots, we established a spectral library of the emission spectra of all six quantum dots for subsequent spectral unmixing analysis. By verifying the linear relationship between the concentration and emission intensity, our system can quantify the concentration of a fluorescent sample from its captured hyperspectral images, which would add an additional dimension of information to qualitative assays. Using linear spectral unmixing, we were able to calculate the proportions of each quantum dot in the homogeneous mixture, and elucidate the spatial distributions of different quantum dots in the heterogeneous mixture, thus allowing for multiple fluorophores to be identified and distinguished from each other. Finally, we have proposed a framework for multiplexed imaging where multiple biomarkers can be simultaneously targeted with different fluorescent probes and imaged in one go, thus paving the way forward for more feasible and highly efficient multiplexed imaging protocols. It provides valuable information for the realization of in vivo fluorescent live imaging with multi-channel viewing.

## Supporting information

Supplemental All

## ASSOCIATED CONTENT

### Supporting Information

Figure S1 shows the full-color reconstructed images of individual quantum dots while they are wet. As compared to Figure 1, all images show a uniform distribution of quantum dots.

**Figure S1.**
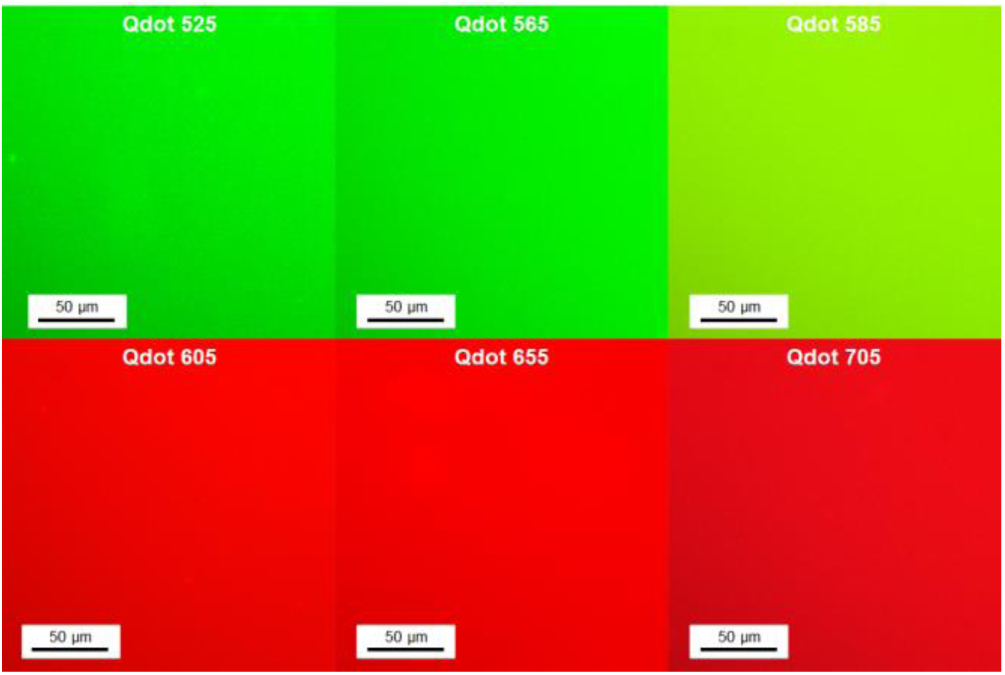
Reconstructed full-color images of individual quantum dots in the wet condition.

Figure S2 shows that we have observed a spectral shift of about 5 nm of the wet and dry samples of all our quantum dot samples.

**Figure S2.**
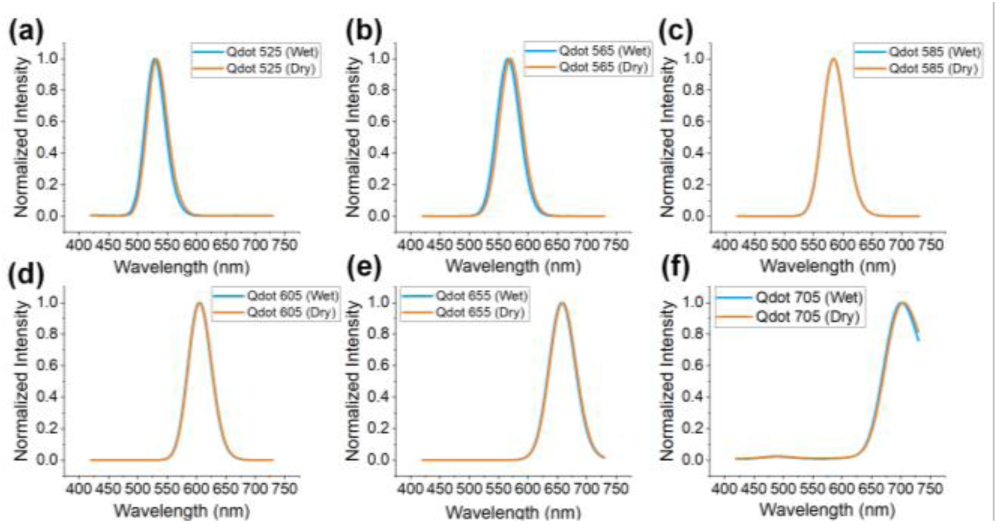
Spectral profiles of all quantum dots in wet and dry conditions.

Another important piece of information that we can derive from the hyperspectral images is the concentration of the quantum dot samples based on their emission intensity. A series of different concentrations of a type of quantum dot sample was imaged. Four solutions of Qdot 605 in different concentrations were prepared at 1 μM (no dilution), 0.75 μM, 0.5 μM, and 0.25 μM respectively, using 1x phosphate-buffered saline (PBS) as the dilution solvent. For each solution, three hyperspectral images of the center of the wet sample were captured, and the average emission spectrum for each solution was calculated and shown in Figure S3(a). The peak intensity values of each solution were then plotted against their concentration values. A linear regression model was fitted to the data with a high *R*^2^ value of 0.90 obtained as seen in Figure S3(b). As such, our system is able to generate a calibration curve for each type of quantum dot sample and subsequently derive its concentration, provided the same excitation conditions are used.

**Figure S3.**
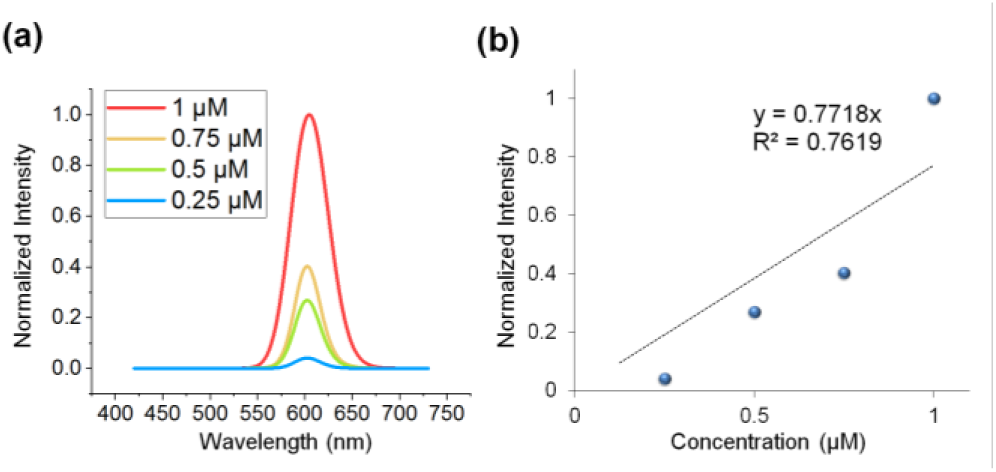
(a) Average emission spectra of different concentrations of Qdot 605. (b) Linear fit on peak intensity values against concentration values of Qdot 605.

The spectral information of the individual quantum dots can be used to predict the spectrum of a homogeneous mixture, and the results are shown in Figure S4. The predicted spectrum of the mixture (“Calculated”) agreed with the measured (“Experimental”) spectrum reasonably well. Discrepancies may be attributed to factors such as the aggregation of quantum dots, and inter-quantum dot energy transfer, which could result in a loss of intensity and spectral shift of the quantum dots.

**Figure S4.**
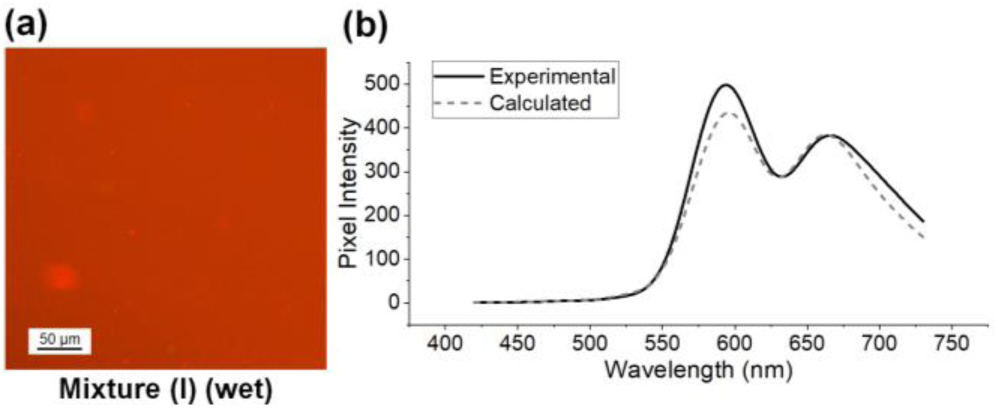
Hyperspectral imaging results of mixture (I). (a) Reconstructed full-color image of the mixture. (b) Calculated and experimentally measured the average spectrum of the mixture within the white area in (a).

## AUTHOR INFORMATION

### Present Addresses

Translational Biophotonics Laboratory, Institute of Bioengineering and Bioimaging, A*STAR, 11 Biopolis Way, Singapore 138667.

## ACKNOWLEDGMENT

The authors would like to acknowledge the funding support from A*STAR Career Development Award (202D800042).

## Notes

### Competing Interest Statement

The authors have declared no competing interest.

