## Supplemental All for "Characterization of Quantum Dots with Hyperspectral Fluorescence Microscopy for Multiplexed Optical Imaging of Biomolecules"

### Supporting Information

Figure S1 shows the full-color reconstructed images of individual quantum dots while they are wet. As compared to Figure 1, all images show a uniform distribution of quantum dots.

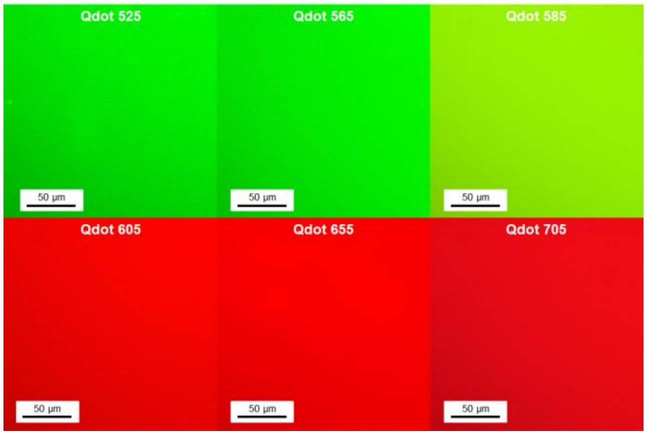


Figure S1. Reconstructed full-color images of individual quantum dots in the wet condition.

Figure S2 shows that we have observed a spectral shift of about 5 nm of the wet and dry samples of all our quantum dot samples.

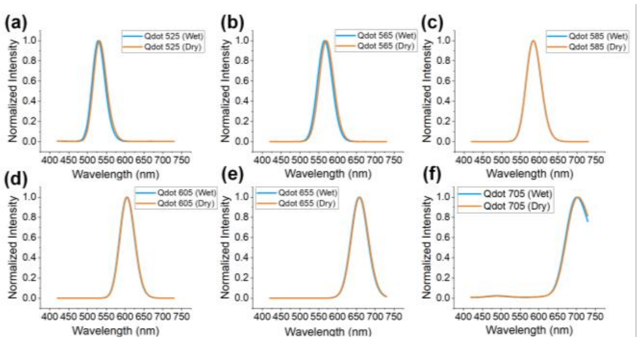


Figure S2. Spectral profiles of all quantum dots in wet and dry conditions.

Another important piece of information that we can derive from the hyperspectral images is the concentration of the quantum dot samples based on their emission intensity. A series of different concentrations of a type of quantum dot sample was imaged. Four solutions of Qdot 605 in different concentrations were prepared at 1 µM (no dilution), 0.75 µM, 0.5 µM, and 0.25 µM respectively, using 1x phosphate-buffered saline (PBS) as the dilution solvent. For each solution, three hyperspectral images of the center of the wet sample were captured, and the average emission spectrum for each solution was calculated and shown in Figure S3(a). The peak intensity values of each solution were then plotted against their concentration values. A linear regression model was fitted to the data with a high $R^2$ value of 0.90 obtained as seen in Figure S3(b). As such, our system is able to generate a calibration curve for each type of quantum dot sample and subsequently derive its concentration, provided the same excitation conditions are used.

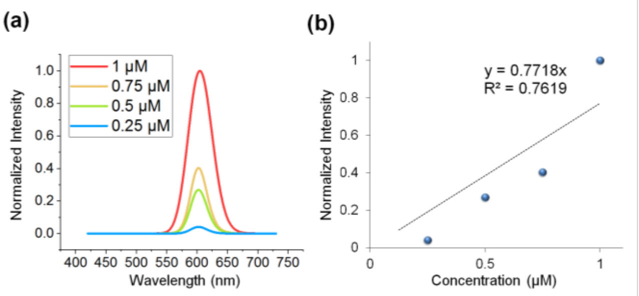


Figure S3. (a) Average emission spectra of different concentrations of Qdot 605. (b) Linear fit on peak intensity values against concentration values of Qdot 605.

The spectral information of the individual quantum dots can be used to predict the spectrum of a homogeneous mixture, and the results are shown in Figure S4. The predicted spectrum of the mixture ("Calculated") agreed with the measured ("Experimental") spectrum reasonably well. Discrepancies may be attributed to factors such as the aggregation of quantum dots, and inter-quantum dot energy transfer, which could result in a loss of intensity and spectral shift of the quantum dots.

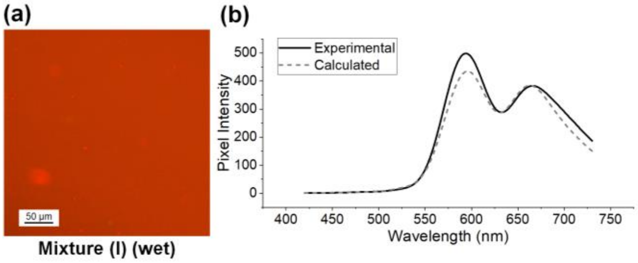


Figure S4. Hyperspectral imaging results of mixture (I). (a) Reconstructed full-color image of the mixture. (b) Calculated and experimentally measured the average spectrum of the mixture within the white area in (a).
